# The molecular basis of antigenic variation among A(H9N2) avian influenza viruses

**DOI:** 10.1101/312967

**Authors:** Thomas P. Peacock, William T. Harvey, Jean-Remy Sadeyen, Richard Reeve, Munir Iqbal

**Affiliations:** Avian Influenza Group, The Pirbright Institute, Pirbright, Woking, UK; Department of Virology, Imperial College London, UK; Boyd Orr Centre for Population and Ecosystem Health, Institute of Biodiversity, Animal Health and Comparative Medicine, College of Medical, Veterinary and Life Sciences, University of Glasgow, UK

**Keywords:** avian influenza, A(H9N2), antigenic evolution, haemagglutinin, receptor-binding avidity, glycosylation, vaccine selection

## Abstract

Avian influenza A(H9N2) viruses are an increasing threat to global poultry production and, through zoonotic infection, to human health where they are considered viruses with pandemic potential. Vaccination of poultry is a key element of disease control in endemic countries, but vaccine effectiveness is persistently challenged by the emergence of antigenic variants. Here we employed a combination of techniques to investigate the genetic basis of H9N2 antigenic variability and evaluate the role of different molecular mechanisms of immune escape. We systematically tested the influence of published H9N2 monoclonal antibody escape mutants on chicken antisera binding, determining that many have no significant effect. Substitutions introducing additional glycosylation sites were a notable exception, though these are relatively rare among circulating viruses. To identify substitutions responsible for antigenic variation in circulating viruses, we performed an integrated meta-analysis of all published H9 haemagglutinin sequences and antigenic data. We validated this statistical analysis experimentally and allocated several new residues to H9N2 antigenic sites providing molecular markers that will help explain vaccine breakdown in the field and inform vaccine selection decisions. We find evidence for the importance of alternative mechanisms of immune escape, beyond simple modulation of epitope structure, with substitutions increasing glycosylation or receptor-binding avidity exhibiting the largest impacts on chicken antisera binding. Of these, meta-analysis indicates avidity regulation to be more relevant to the evolution of circulating viruses, suggesting that a specific focus on avidity regulation is required to fully understand the molecular basis of immune escape by influenza, and potentially other viruses.

## Introduction

In recent years novel avian influenza virus (AIV) strains have emerged as a major threat to animal and human health. H9N2 AIVs are endemic across much of Asia, the Middle East, and North Africa, where they cause severe economic losses to the poultry industry through moderate-to-high morbidity and mortality^1–3^. H9N2 viruses are an emerging threat to poultry in new geographic regions, with the first reported sequences found in Russia, Sub-Saharan Africa, and Indonesia occurring within the last year^4–7^. Additionally, certain H9N2 lineages are considered to have pandemic potential due to their repeated isolation from humans and their ability to adapt to and transmit between ferrets, the animal model for human transmission^8–10^. Certain H9N2 virus internal gene constellations have a unique capacity to elevate the zoonotic potential of non-H9N2 AIVs with recent examples including H7N9, H10N8, and clade 2.3.4.4 H5N6, all known to have high mortality rates in humans^11^.

Due to the threat to poultry and human health posed by endemic and emerging H9N2 AIVs, many countries vaccinate poultry as a major method of viral control, with conventional inactivated vaccines being used most commonly^12–14^. However, as with human influenza virus, poor vaccine matching due to antigenic drift often results in vaccine failure^3,12,14,15^. Immunity, following infection or vaccination, is primarily achieved by the generation of neutralizing antibodies that interact with haemagglutinin (HA), the major influenza antigen and receptor-binding protein, sterically blocking attachment to target cells^16^. Consequently, the antigenic variability of influenza viruses, commonly assessed using the haemagglutination inhibition (HI) assay, can largely be mapped to amino acid substitutions near the HA receptor-binding site^17,18^. These substitutions have been found to contribute to both actual and apparent antigenic change via a variety of mechanisms, namely changes to epitope structure, acquisition of additional glycosylation sites, and modulation of receptor-binding avidity^19–21^.

Amino acid substitutions that alter the biophysical properties (shape, charge, polarity, etc.) of an epitope have the potential to cause antigenic change by directly affecting antibody binding. This is the most conventional mechanism of immune escape. The acquisition of additional N-linked glycosylation sites is another mechanism by which influenza viruses may escape antibody neutralization^20,21^; bulky oligosaccharides can sterically ‘shield’ HA epitopes from antibody recognition and the antibody response is potentially directed to alternative antigenic sites^22^. In the absence of compensatory mutations, additional glycosylation has been described to reduce receptor-binding avidity, which has been hypothesized to restrict hyper-glycosylation as an immune evasion strategy^21,23^. Neutralizing antibodies competitively inhibit HA-receptor interactions. Amino acid substitutions that modulate receptor-binding avidity can therefore contribute to apparent antigenic change detected by HI assays. Modulation of avidity has been hypothesized to be a true form of immune escape, rather than an artefact of the HI assay^19,24^. The relative roles of each of these mechanisms in evolution across influenza subtypes and the consequences for vaccine effectiveness is not well known.

Previously, we used a ‘classical’ monoclonal antibody (mAb) escape mutant approach to investigate the antigenic architecture of the H9 HA, identifying several residues that were assigned to two discrete antigenic sites, ‘H9-A’ and ‘H9-B’^25^. Although these residues were important for the binding of murine mAbs, their significance in the context of circulating field viruses remained unknown. Many residues identified in our previous study, as well as several similar published studies^26–30^, are completely conserved amongst circulating viruses indicating they are not responsible for antigenic variability observed in the field.

In this paper, we identify amino acid residues and substitutions responsible for H9N2 virus antigenic variability in the field through a variety of approaches. We reconstitute mutations at every published H9 HA monoclonal escape mutant position^25–30^ and test the effect of these substitutions on polyclonal chicken antisera binding. We then adapt an approach previously developed for human influenza viruses^17^, performing a meta-analysis of all published H9 HI assay and corresponding genetic data to identify naturally-occurring antigenically-important substitutions. We identify substitutions that correlate with antigenic changes and prove causation using reverse genetics and HI assays. Several novel antigenic residues are predicted, which we demonstrate are indeed relevant to the antigenic diversity of field H9N2 viruses. Overall, we provide a comprehensive and systematic analysis of the molecular basis of H9 HA antigenic diversity in chickens, which are routinely vaccinated by the poultry industry and hence the critical host species for understanding risks of vaccine escape.

## Results

### Antigenic impact of escape mutant viruses

Six studies describe a total of 39 unique mAb escape mutations in H9N2 HA1 across 30 amino acid positions^25–30^ (Table S1). Viruses were generated with mutations at each published escape residue in the HA of the H9N2 virus A/chicken/Pakistan/UDL-01/2008 (UDL1/08) and tested using HI, with a panel of 8 anti-UDL1/08 chicken post-infection antisera, to assess the impact of these mutations on antigenicity. When multiple escape mutants were reported at the same position, the mutation, or mutations, that introduced the largest difference in biophysical properties were generated (e.g. of T129K and T129A, only T129K was tested). Furthermore, alternative, more biologically relevant mutations at escape sites were introduced in some cases (64% of published escape mutations are absent or almost absent (frequency <1%) amongst sequenced H9N2 viruses, Table S2). Mutants unable to be rescued in the UDL1/08 background were dropped from further study (Table S3).

In total, 44 mutants at 30 positions described in escape mutant studies were assessed by HI assay (Fig. 1). Of these, 25 at 20 positions showed significant reductions in HI titre compared to wild type UDL1/08. Several of the introduced substitutions – 19 at 16 amino acid positions – caused no significant change in titre. Interestingly, of the seven substitutions with the largest impact on chicken antiserum binding (≥1.30 log_2_ titre), four corresponded to the addition of glycosylation sites (T127N, L150S, T188N, and D189N), while the remaining three corresponded to known receptor-binding residues (T145I, L216Q and I217Q)^9,31–33^. Each of these four glycosylation mutants were tested for the presence of the additional glycans by western blot and all displayed upward band shifts, consistent with the addition of glycosylation sites (Fig. S1). Complete N-linked glycan motifs at the sites of the four glycosylation site mutants tested are, however, rare among sequenced field isolates (Table 1).

**Fig 1.**
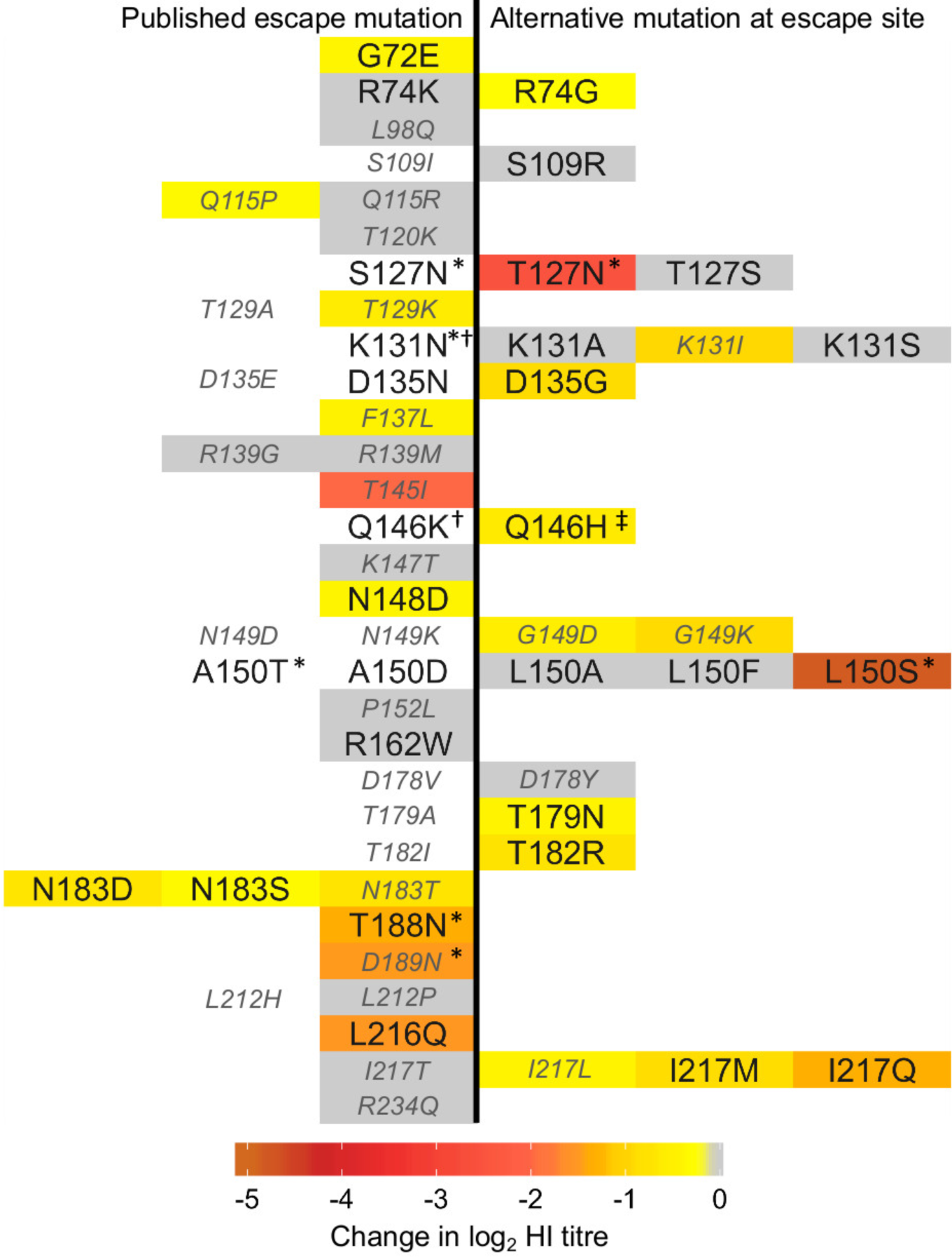
Impact of mutation at published mAb escape sites on haemagglutination inhibition titres performed with polyclonal chicken antisera. Change in log_2_ HI titre associated with stated amino acid replacements, based on 8 individual anti-UDL1/08 antisera with a minimum of 5 technical repeats, are indicated by cell colouring according to the colour legend with non-significant differences shown in grey and untested substitutions in white. Published mAb escape mutations are shown to the left of centre while alternative substitutions are shown to the right. Mutations that introduce absent or almost absent amino acid states (<1% of sequenced natural isolates) are labelled in smaller, italicized, grey font. * addition of potential glycosylation site. ^†^ would not rescue. ^‡^ Q146H rescued with additional substitution T186K.

**Table 1.** Frequency of potentially antigenic glycosylation sites in natural H9 isolates.

| Potential glycosylation site position<sup>a</sup> | Mutant(s) in this study<sup>b</sup> | Frequency<sup>c</sup> |
| --- | --- | --- |
| 127-129 | S127N, **T127N** | 6.8% |
| 131-133 | K131N | 0.1% |
| 148-150 | A150T, **L150S** | 1.5% |
| 188-190 | **T188N** | 2.6 |
| 189-191 | **D189N** | 0.1% |

<sup>a</sup>Potential glycosylation sites defined as amino acid motif N-X-S/T, where X is not P;
<sup>b</sup>Mutants described in Fig. 1, mutants made in this study shown in bold; <sup>c</sup>Total frequency of potential glycosylation site at this position, based on full length H9 sequences in the NCBI database as of July 2017.

Of the tested mutations found to significantly affect chicken antiserum binding, several are unable to contribute significantly to antigenic variability among circulating strains due to their rareness in nature. For example, positions 115, 129, 145, and 189 (as well as non-significant positions 98, 137, 147, 152, 212, and 234) are all >98% conserved (Table S2). For a sample of viruses covering all major H9N2 lineages, amino acid identity at each of the 20 escape mutant sites where a significant impact on HI titre was detected is shown in Fig. 2 (an expanded figure including all 30 escape mutant sites is shown in Fig. S2A). Of the 39 published escape mutations, 27 (69%) had no significant impact on HI titres, were absent or almost absent among sequenced viruses, or both (Fig. 1). At 10 of the 30 published escape sites, no evidence of an antigenic impact in HI assays was detected, even when taking alternative substitutions at the site into account. These results suggest that antibodies in chicken polyclonal antisera do not consistently recognise the same epitopes as mouse mAbs or that there are differences in immunodominance, and that further sites likely contribute to antigenic variation of H9N2 viruses in nature.

**Fig 2.**
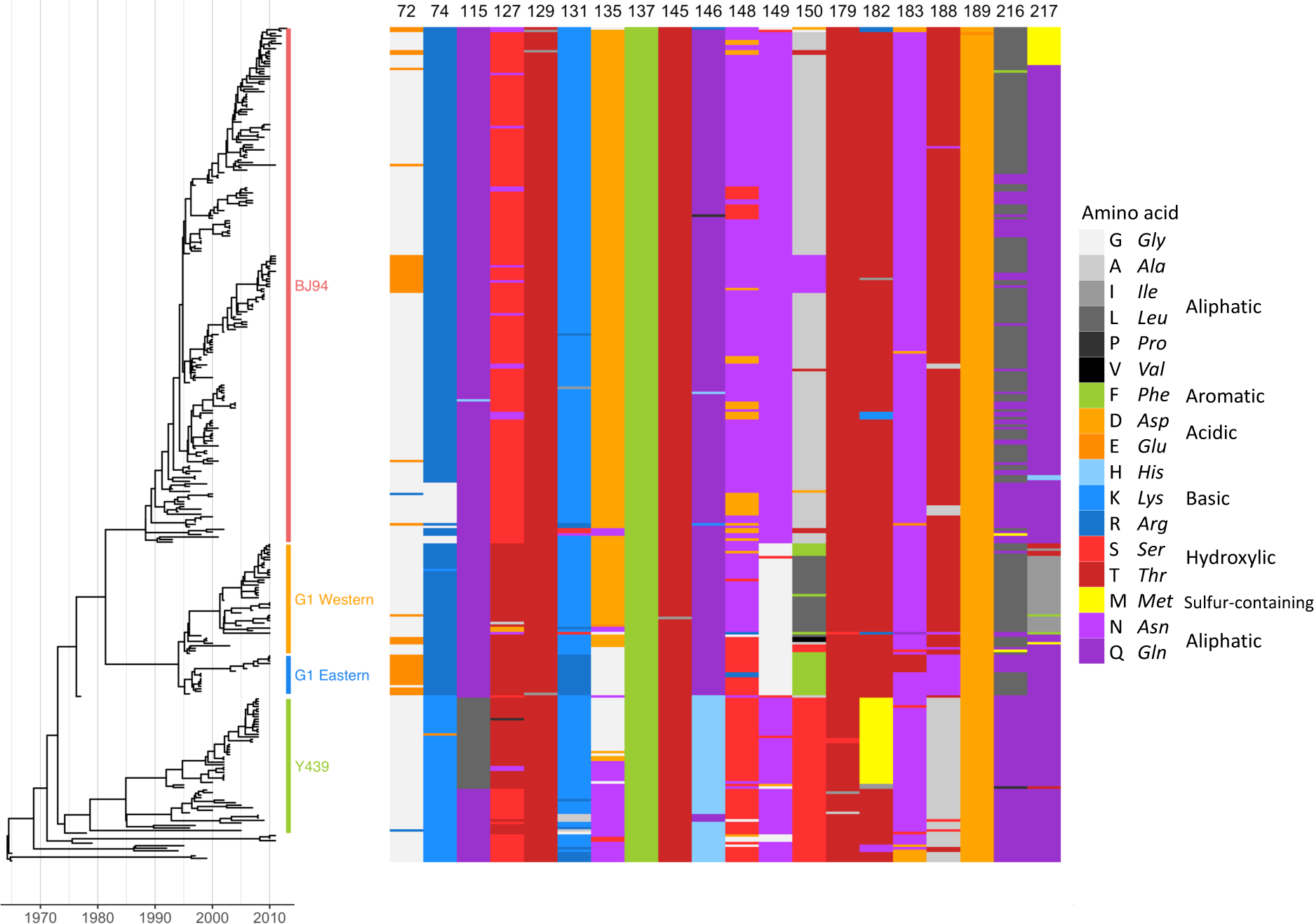
Sequence variation at antigenically significant H9 haemagglutinin escape mutant sites. Time-resolved Bayesian HA1 phylogeny and amino acid identity at each of the 20 mAb escape residues found to significantly affect HI titres with polyclonal chicken antisera. Amino acid identity is shown by colour, grouped by side-chain property, according to the legend. Each virus (n = 330) included in the phylogeny has associated HI data and was included in integrated modelling of genetic and antigenic data.

### Modelling of HI and genetic data and prediction of antigenically pertinent substitutions

To investigate genetic differences responsible for H9N2 field strain antigenic diversity, we performed an integrated statistical analysis of antigenic and genetic data. All published HI data for H9N2 with associated HA1 nucleotide sequences were compiled (citations in Table S4) and supplemented with additional HI assays (Table S5). The assembled dataset comprised 330 H9N2 virus sequences covering all major lineages (Fig. 2), including 103 viruses against which antisera were raised, with a total of 2,574 individual HI titres measured between 2,131 unique combinations of virus and antiserum.

Linear mixed models were used to account for variation in HI in terms of viral sequence changes, as previously described^17^. Initial model selection determined variables representing systematic differences in the magnitudes of titres recorded for individual test viruses, for antisera raised against particular reference viruses, for different serum types (i.e. live virus vs inactivated vaccination, and chicken vs ferret), and for different studies. To control for repeated measures that arise from phylogenetic correlations in the data and prevent false positive support for substitutions, branches of the HA1 phylogeny correlated with drops in HI titre were identified next, and then terms representing substitutions at each variable HA1 position were tested to determine whether their inclusion significantly improved model fit.

This analysis identified antigenically distinct amino acids at 12 HA1 positions (74, 121, 131, 135, 150, 180, 183, 195, 198, 216, 249, and 288). Of these, seven had been identified in published H9N2 mouse mAb escape studies (74, 131, 135, 150, 183, 188, and 216), including positions in previously described antigenic sites (Table 2)^25,27^. Several additional residues were identified when ferret or chicken antisera datasets were analysed independently and were also considered (72, 217, 264, 276, and 306), a further two (72 and 217) had also been described by escape mutant studies^25,26^. Among the specific substitutions inferred to be antigenically relevant, G72E, K131N, N183T, and L216Q are published mAb escape mutations; G72E, N183T, and L216Q (K131N could not be rescued) were also confirmed to significantly reduce binding by polyclonal chicken antisera in HI assays above (Fig. 1). Amino acid identity at each of the 17 HA positions identified using this modelling approach is shown for each virus included in the analysis in Fig. S2B.

**Table 2.**
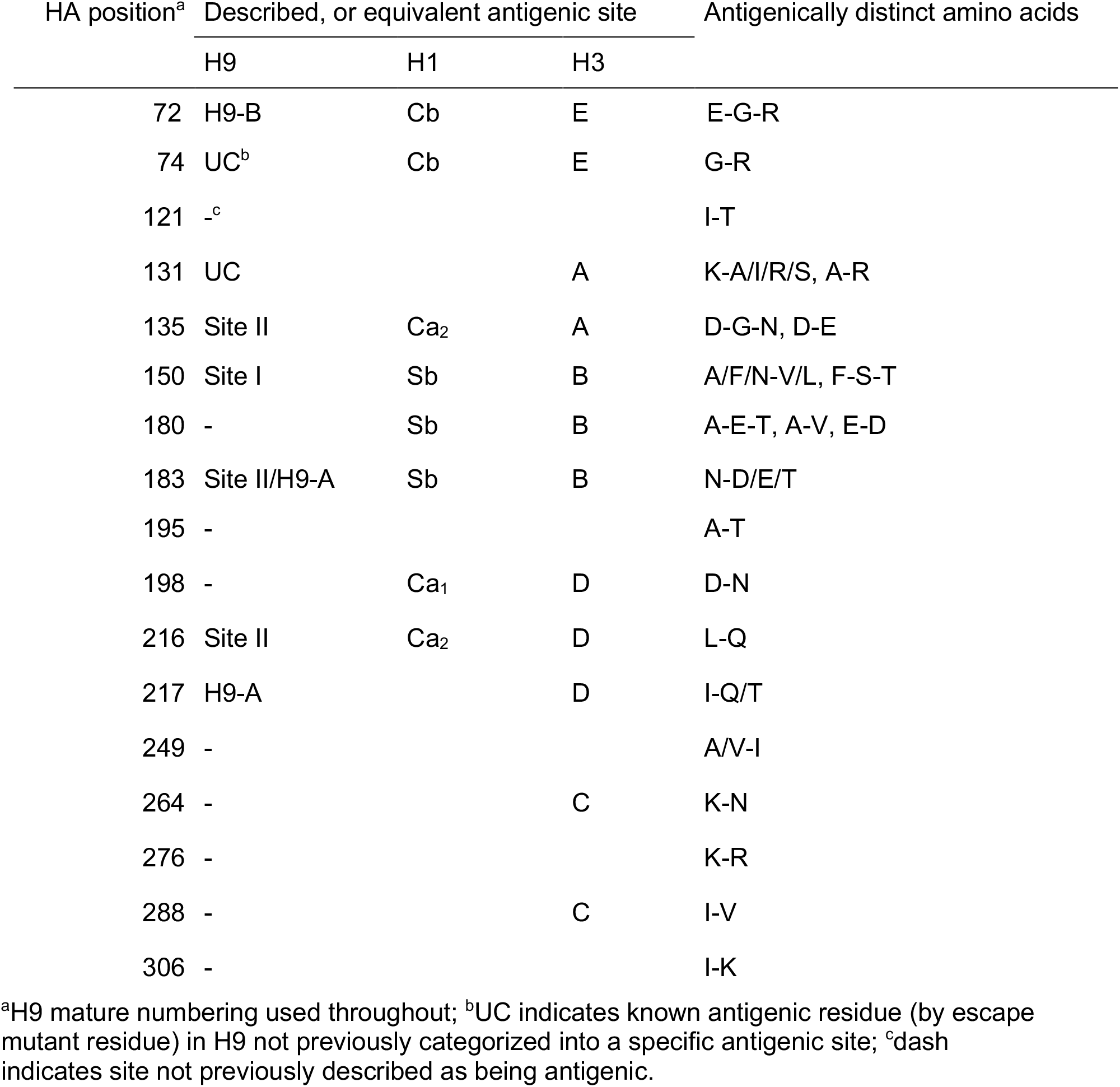
Amino acids substitutions correlating with antigenic change and corresponding antigenic sites.

| HA position<sup>a</sup> | Described, or equivalent antigenic site |  |  | Antigenically distinct amino acids |
| --- | --- | --- | --- | --- |
|  | H9 | H1 | H3 |  |
| 72 | H9-B | Cb | E | E-G-R |
| 74 | UC<sup>b</sup> | Cb | E | G-R |
| 121 | -<sup>c</sup> |  |  | I-T |
| 131 | UC |  | A | K-A/I/R/S, A-R |
| 135 | Site II | Ca<sub>2</sub> | A | D-G-N, D-E |
| 150 | Site I | Sb | B | A/F/N-V/L, F-S-T |
| 180 | - | Sb | B | A-E-T, A-V, E-D |
| 183 | Site II/H9-A | Sb | B | N-D/E/T |
| 195 | - |  |  | A-T |
| 198 | - | Ca<sub>1</sub> | D | D-N |
| 216 | Site II | Ca<sub>2</sub> | D | L-Q |
| 217 | H9-A |  | D | I-Q/T |
| 249 | - |  |  | A/V-I |
| 264 | - |  | C | K-N |
| 276 | - |  |  | K-R |
| 288 | - |  | C | I-V |
| 306 | - |  |  | I-K |

<sup>a</sup>H9 mature numbering used throughout; <sup>b</sup>UC indicates known antigenic residue (by escape mutant residue) in H9 not previously categorized into a specific antigenic site; <sup>c</sup>dash indicates site not previously described as being antigenic.

### Antigenic characterization of mutant viruses to test predictions

To validate the predictions of modelling HI and genetic data, we tested mutant viruses using HI assays and compared titres to those measured using parental viruses. Mutant libraries were generated in the backgrounds of three antigenically distinct H9N2 viruses, UDL1/08, A/Hong Kong/33982/2009 (HK/33982) and A/chicken/Emirates/R66/2002 (Em/R66). Wherever possible, mutations were made that were reciprocal between these viruses (i.e. UDL1/08 naturally has 183N and HK/33982 has 183T, substitution between N and T at 183 is predicted to have an antigenic effect therefore UDL1/08 N183T and HK/33982 T183N were generated and tested). When reciprocal substitutions were introduced, both mutant viruses were tested using antiserum raised against both parent viruses. Changes in mean log_2_ titre relative to the parent virus, against both parental and mutant-like antiserum, are detailed in Table S6.

Of the 26 amino acid substitutions from Table 2 introduced, 19 resulted in significantly different titres measured when using parental antiserum (G72E, E72G, R74G, I121T, T121I, K131I, D135G, A150L, A180E, E180A, N183D, N183T, T183N, L216Q, Q216L, I217Q, Q217I, I249V and V249I), while seven did not (K131A, K131S, G135D, L150A, L150F, F150L and D198N). For substitutions introduced into the UDL1/08 background at escape mutation sites, effect sizes are shown in Fig. 1 and where reciprocal forward and reverse mutations were introduced in different backgrounds, effect sizes are shown in Fig. 3. In Fig. 3, lower or higher HI titres recorded for mutant viruses, relative to the parental virus, are shown in shades of orange and blue respectively. Generally, drops in titres were observed against the parental antiserum and increases were observed against antiserum possessing the introduced substitution, suggesting reduced and increased antibody recognition respectively (Fig. 3).

**Fig 3.**
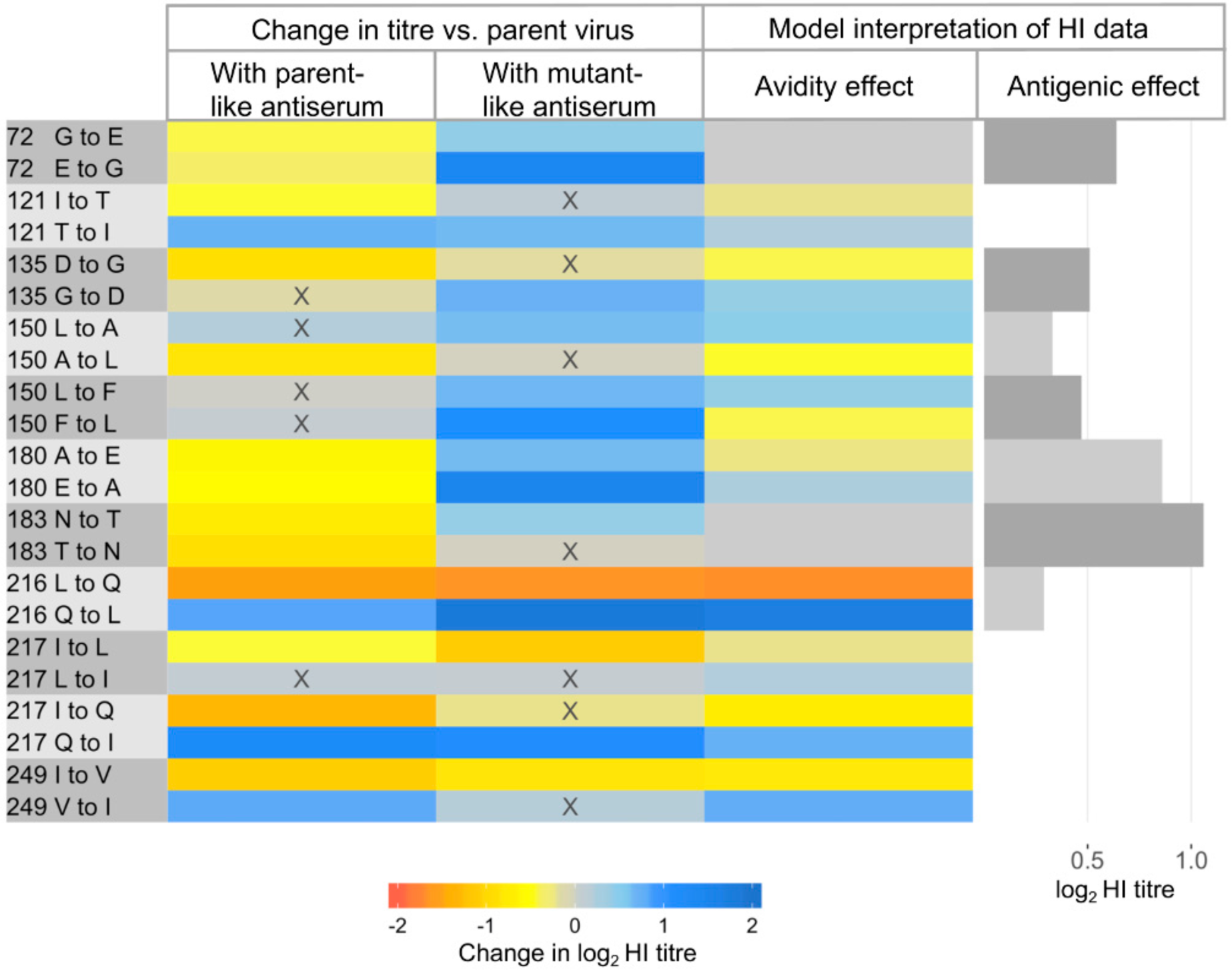
Heat-map showing the impacts of reciprocal amino acid substitutions introduced at sites identified by modelling. Changes in HI titre (log_2_) associated with each introduced substitution are shown by heat map, measured using antiserum both lacking the introduced substitution (parent-like) and possessing the introduced mutation (mutant-like) from combinations of the viruses UDL1/08, Em/R66 and HK/33982. Changes in HI titre associated with each pair of substitutions were interpreted as effects of altered avidity and antigenicity using Equation 1, are also shown. Avidity effects cause an increase (blue) or drop (orange) in titre depending on the direction of substitution while antigenic effects are the estimated change in titre resulting from antigenic dissimilarity. X indicates not individually significant.

Testing mutant viruses using antisera raised both against the parent virus, and against a virus possessing the introduced amino acid state, allowed us to calculate the relative contributions of antigenic and avidity effects (Equation 1, Materials and Methods). The impact on HI titre of altered antigenicity and avidity associated with each pair of reciprocal substitutions is shown in Fig. 3 (and Table S6). Of the 11 reciprocal pairs considered, two were estimated to influence HI titres as a result of altered antigenicity alone (72G/E and 183N/T), four as a result of altered avidity only (121I/T, 217L/Q, 217I/Q and 249I/V) and five as a combination of these effects (135D/G, 150A/L, 150F/L, 180A/E and 216L/Q) indicating that both these mechanisms of immune escape are potentially playing a role in the evolution of H9N2 viruses.

The impact of substitutions between A and E at position 180 show a classic antigenic signature (Fig. 3). Both A180E and E180A were associated with significant drops in log_2_ titre of 0.61 and 0.54 measured using antiserum raised to the parental virus, UDL1/08 (180A) and Em/R66 (180E) respectively. Both A180E and E180A also increased titres, by 0.65 and 1.51 respectively, measured using mutant-like antiserum that possessed the introduced amino acid at position 180. These changes in titre indicate that the A180E substitution has decreased and increased the reactivity of antibodies raised against 180A and 180E viruses, respectively; a consistent pattern was observed with the E180A substitution. Consequently, a sizeable antigenic effect (0.86 log_2_ titre) is estimated alongside a smaller avidity effect (+/− 0.25 log_2_ titre). Similar patterns of change in titres indicated sizeable antigenic effects for substitution between G and E at position 72 and between N and T at 183.

Exchange of Q and L at position 216 represents an example of a substitution impacting HI titres due to altered receptor-binding avidity (Fig. 3). L216Q reduced titres against antisera both lacking the introduced amino acid state, UDL1/08 (216L) and possessing it, Em/R66 (216Q). Conversely, Q216L increased titres against antisera both lacking (Em/R66) and possessing (UDL1/08) the introduced amino acid state. These changes indicate that an H9 HA possessing 216Q has higher avidity than a virus with 216L, resulting in lower HI titres generally, regardless of whether antiserum was raised to a 216Q or 216L virus. We were able to confirm this conclusion in a receptor destroying enzyme-based avidity assay, where swapping the single amino acid at position 216 between UDL1/08 and Em/R66 allowed an almost complete swap of avidity phenotypes (Fig. S3). Similar effects of avidity on HI titres were observed at positions 121, 217 and 249. Interestingly, L216Q, I217L and I217Q, which each result in drops in HI titre due to altered avidity (Fig. 3) are either published mAb escape mutations or occur at the same position as escape mutations (Fig. 1). This indicates that increased avidity facilitated by single amino acid substitutions is a mechanism of immune escape for H9.

Closer observation of the pattern of change in HI titre at position 135 suggests an asymmetric antigenic effect of substitutions between D and G at this position. The average impacts on titre of either D135G or G135D were significant and large (−0.90 and +0.73) when measured with antisera raised against 135D containing UDL1/08. In contrast, no significant impact on titres were observed using antisera raised against 135G containing HK/33982, consistent with 135D belonging to an immunodominant epitope. A model that estimated two antigenic effects of substitutions between D and G, depending on which antiserum was used, rather than a single antigenic effect was found to improve model fit as assessed by deviance information criterion (DIC). Experimental studies have previously demonstrated an asymmetric effect at the homologous HA position in human A(H3N2) viruses^24^. The same asymmetric model was also favoured in an analysis of H3 HI titres associated with the asymmetric effect reported by Li *et al*. (2013)^24^.

### Mapping of substitutions into antigenic sites using mAbs

Previously, we described a panel of mouse mAbs against UDL1/08 binding to two discrete antigenic sites, ‘H9-A’ and ‘H9-B’^25^. These mAbs were used to perform HI assays against every mutant virus generated in this study. If a mutant virus had lower titres to mAbs from either group, but not the other, the residue was assigned to the corresponding antigenic site. The mutants G149K, T179N, A180D and A180E reduced titres to H9-A binding mAbs indicating positions 149, 179 and 180 likely form part of antigenic site H9-A alongside residues 145, 146, 183, 212, 217, and 234. Titres for mutants G72E and D135G indicated these positions form part of site H9-B alongside residues 115, 120, 139 and 162 (Fig. 4, Tables 3 and S7). The four glycosylation mutants (T127N, L150S, T188N, and D189N) reduced titres to both H9-A and H9-B mAbs suggesting that the addition of bulky glycans may block antibody binding to both sites. The substitutions K131I, D178Y, T182R, L216Q, I217M and I217Q also slightly reduced binding of mAbs from both groups. These residues may either lie between antigenic sites and exert an effect on both, or, these substitutions may overcome antibody binding by increasing HA receptor-binding avidity.

**Fig 4.**
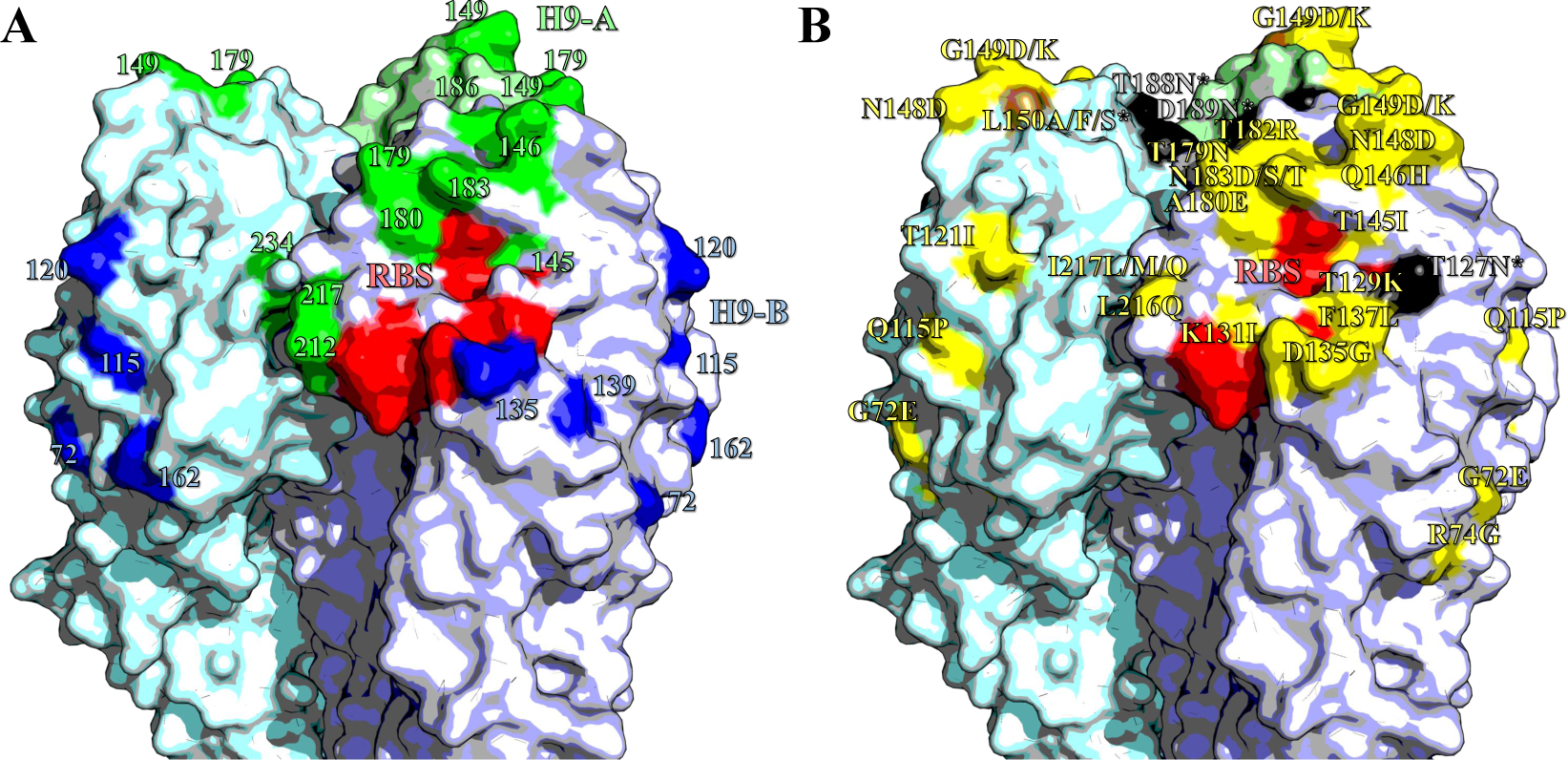
Antigenic structure of H9 HA. Homotrimers of H9 HA. Selected receptor-binding residues shown in red (P92, G128, T129, S130, S131, A132, W142, N173, L184, Y185, N214, G215, G218 and R219). Images made in Pymol^52^ (Schrödinger) based on the structure of A/swine/Hong Kong/9/1998 (Protein databank ID:1JSD)^53^. (A) Residues recognized by mouse mAbs, positions with updated site H9-A shown in green, H9-B residues shown in blue. (B) Residues labelled with substitutions that affect the binding of chicken polyclonal antisera. Non-glycosylation altering substitutions shown in yellow; glycosylation site adding mutations shown in black; site 150, which had both glycosylation adding and non-glycosylation adding mutations shown in brown.

**Table 3.**
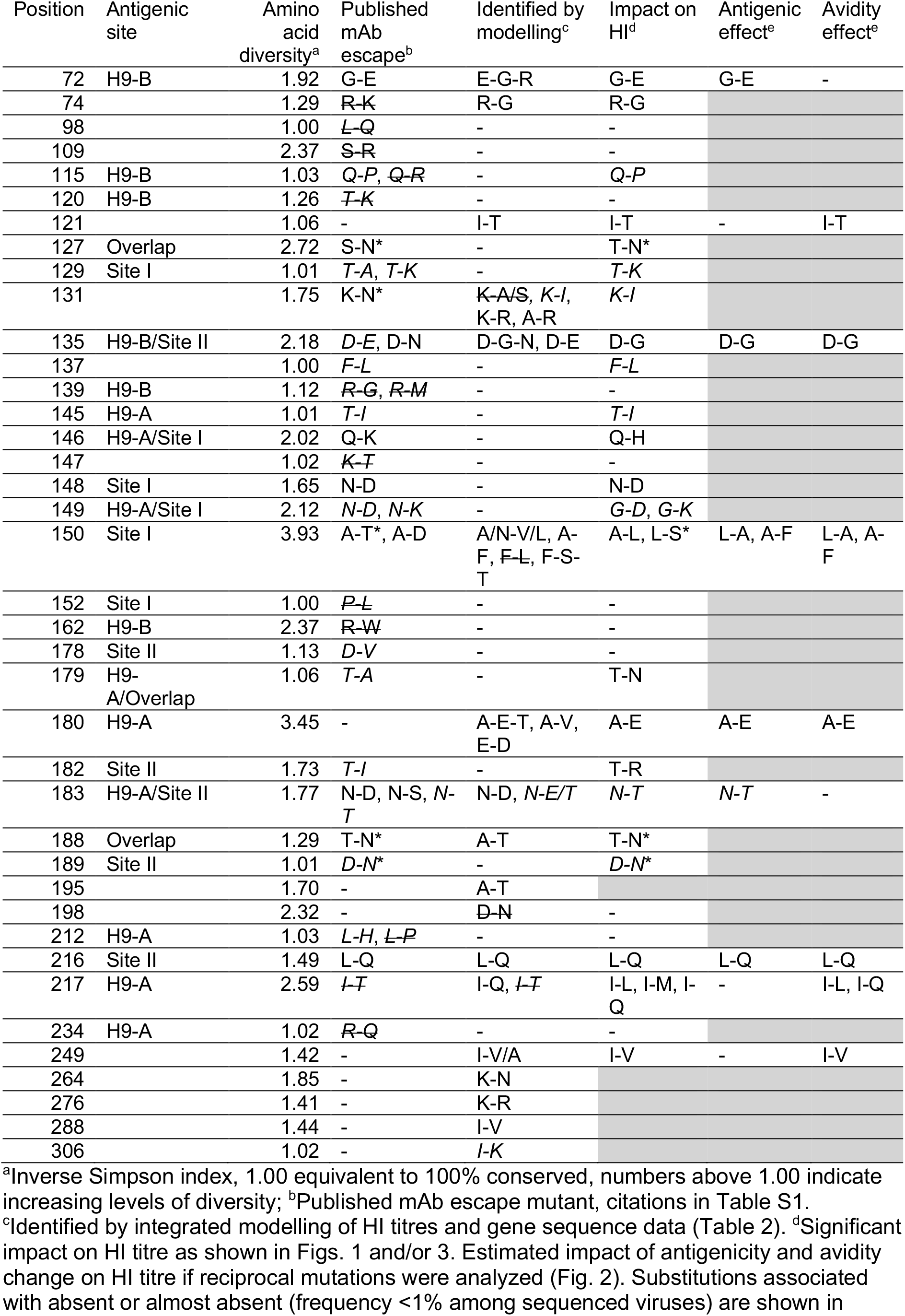
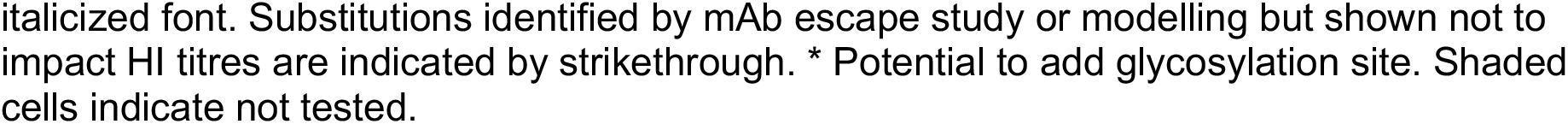
Summary of residues investigated in this study and their relation to antigenicity and avidity.

| Position | Antigenic site | Amino acid diversity<sup>a</sup> | Published mAb escape<sup>b</sup> | Identified by modelling<sup>c</sup> | Impact on HI<sup>d</sup> | Antigenic effect<sup>e</sup> | Avidity effect<sup>e</sup> |
| --- | --- | --- | --- | --- | --- | --- | --- |
| 72 | H9-B | 1.92 | G-E | E-G-R | G-E | G-E | - |
| 74 |  | 1.29 | ~~R-K~~ | R-G | R-G |  |  |
| 98 |  | 1.00 | ~~L-Q~~ | - | - |  |  |
| 109 |  | 2.37 | ~~S-R~~ | - | - |  |  |
| 115 | H9-B | 1.03 | Q-P, ~~Q-R~~ | - | Q-P |  |  |
| 120 | H9-B | 1.26 | ~~T-K~~ | - | - |  |  |
| 121 |  | 1.06 | - | I-T | I-T | - | I-T |
| 127 | Overlap | 2.72 | S-N* | - | T-N* |  |  |
| 129 | Site I | 1.01 | T-A, T-K | - | T-K |  |  |
| 131 |  | 1.75 | K-N* | ~~K-A/S~~, K-I, K-R, A-R | K-I |  |  |
| 135 | H9-B/Site II | 2.18 | D-E, D-N | D-G-N, D-E | D-G | D-G | D-G |
| 137 |  | 1.00 | F-L | - | F-L |  |  |
| 139 | H9-B | 1.12 | ~~R-G~~, ~~R-M~~ | - | - |  |  |
| 145 | H9-A | 1.01 | T-I | - | T-I |  |  |
| 146 | H9-A/Site I | 2.02 | Q-K | - | Q-H |  |  |
| 147 |  | 1.02 | ~~K-T~~ | - | - |  |  |
| 148 | Site I | 1.65 | N-D | - | N-D |  |  |
| 149 | H9-A/Site I | 2.12 | N-D, N-K | - | G-D, G-K |  |  |
| 150 | Site I | 3.93 | A-T*, A-D | A/N-V/L, A-F, ~~F-L~~, F-S-T | A-L, L-S* | L-A, A-F | L-A, A-F |
| 152 | Site I | 1.00 | ~~P-L~~ | - | - |  |  |
| 162 | H9-B | 2.37 | ~~R-W~~ | - | - |  |  |
| 178 | Site II | 1.13 | D-V | - | - |  |  |
| 179 | H9-A/Overlap | 1.06 | T-A | - | T-N |  |  |
| 180 | H9-A | 3.45 | - | A-E-T, A-V, E-D | A-E | A-E | A-E |
| 182 | Site II | 1.73 | T-I | - | T-R |  |  |
| 183 | H9-A/Site II | 1.77 | N-D, N-S, N-T | N-D, N-E/T | N-T | N-T | - |
| 188 | Overlap | 1.29 | T-N* | A-T | T-N* |  |  |
| 189 | Site II | 1.01 | D-N* | - | D-N* |  |  |
| 195 |  | 1.70 | - | A-T |  |  |  |
| 198 |  | 2.32 | - | ~~D-N~~ | - |  |  |
| 212 | H9-A | 1.03 | L-H, ~~L-P~~ | - | - |  |  |
| 216 | Site II | 1.49 | L-Q | L-Q | L-Q | L-Q | L-Q |
| 217 | H9-A | 2.59 | ~~I-T~~ | I-Q, ~~I-T~~ | I-L, I-M, I-Q | - | I-L, I-Q |
| 234 | H9-A | 1.02 | ~~R-Q~~ | - | - |  |  |
| 249 |  | 1.42 | - | I-V/A | I-V | - | I-V |
| 264 |  | 1.85 | - | K-N |  |  |  |
| 276 |  | 1.41 | - | K-R |  |  |  |
| 288 |  | 1.44 | - | I-V |  |  |  |
| 306 |  | 1.02 | - | I-K |  |  |  |

<sup>a</sup>Inverse Simpson index, 1.00 equivalent to 100% conserved, numbers above 1.00 indicate increasing levels of diversity; <sup>b</sup>Published mAb escape mutant, citations in Table S1.
<sup>c</sup>Identified by integrated modelling of HI titres and gene sequence data (Table 2). <sup>d</sup>Significant impact on HI titre as shown in Figs. 1 and/or 3. Estimated impact of antigenicity and avidity change on HI titre if reciprocal mutations were analyzed (Fig. 2). Substitutions associated with absent or almost absent (frequency <1% among sequenced viruses) are shown in
italicized font. Substitutions identified by mAb escape study or modelling but shown not to impact HI titres are indicated by strikethrough. \* Potential to add glycosylation site. Shaded cells indicate not tested.

## Discussion

Determining the molecular mechanisms of antigenic evolution is vital to understanding viral disease dynamics, and vaccine selection and development. We have presented here a systematic analysis of the molecular basis of H9N2 antigenicity. We assessed the contribution of residues reported by the literature to be antigenic and their relation to virus antigenic diversity and showed that previous studies based on murine monoclonal antibody escape mutants are only moderately reliable at providing relevant antigenic information; several false positives found to have no impact on inhibition of chicken antisera were identified, as were residues that are very highly conserved in the field. Furthermore, we carried out a meta-analysis of all available matching genetic and antigenic data for H9N2 viruses, modelling the relationship between genotype and antigenic phenotype to identify novel residues directly responsible for observed antigenic diversity in the field, and we categorized these residues into recently defined antigenic sites. Analysis of reciprocal mutations introduced by reverse genetics allowed us to differentiate multiple mechanisms facilitating immune escape including addition of glycans and increased receptor-binding avidity, as well as conventional changes to epitope structure that directly affect antibody binding.

In this study substitutions with the largest impact of antigenicity were associated with two distinct mechanisms of immune evasion. Of the eight most significant mutants, four (T127N, L150S, T188N and D189N) possessed additional glycosylation sites and four (T145I, L216Q, I217Q, and I249V) possessed substitutions either known to influence receptor-binding avidity in H9N2 viruses or determined to do so in this study through analysis of reciprocal mutants, or both^9,31–33,35^. These observations demonstrate the potential of alternative mechanisms, other than reduction of antibody recognition by altering epitope biophysics, in facilitating immune escape and determining antigenic diversity of H9N2 viruses. Substitutions altering avidity were well represented among those identified by modelling of antigenic and genetic data collected from H9N2 field strains suggesting that avidity regulation may be an important mechanism of immune escape in nature. In contrast, substitutions associated with acquisition of additional glycans were largely absent suggesting this to be a less important mechanism, possibly due to an associated fitness cost, as reported elsewhere for human influenza^21^.

Implementing a systematic approach allows for the comprehensive evaluation of the phenotypic impact of amino acid substitutions. For example, we identify substitutions at the novel antigenic residue 180 that explain significant antigenic variation among natural viruses sampled across the H9N2 phylogeny. The analysis of reciprocal substitutions can then differentiate between mechanisms of immune escape, exchange between A and E at 180 was indicated to influence HI titres due to altered antibody-epitope recognition rather than modulation of receptor-binding avidity. Epitope mapping using mAbs further indicated that residue 180 is a constituent of the H9-A antigenic site.

We were able to detect a likely example of changing immunodominance, demonstrating that such effects can be distinguished using HI data. Substitutions between aspartic acid and glycine at position 135 were observed to impact titres measured using antisera raised to a D135 virus but not using antisera raised to a G135 virus. These results suggest that an epitope involving position 135 is immunodominant when a virus possesses D135 (aspartate has a prominent acidic side chain), but that antibody responses generated against viruses possessing G135 are directed to other regions (unsurprising, given that glycine is the smallest amino acid). Substantial change in immunodominance resulting from a single amino acid substitution has previously been reported in human influenza^24^. While we did not detect any other cases of asymmetric antigenic change, we speculate that the phenomenon is perhaps widespread though hard to detect, except in the most dramatic cases. Better understanding of this phenomenon may facilitate the development of vaccines designed to guide the immune response towards more conserved epitopes providing better cross-protection.

In contrast to previously validated applications of the described modelling technique to human influenza A(H1N1) and foot-and-mouth disease virus where very large antigenic datasets (>10,000 titres) were collected by a single laboratory under relatively consistent methods^17,36^, the data analysed here were drawn from multiple sources and encompass varying experimental designs, serum types, and production methods. This is expected to limit the accuracy of parameter coefficient estimates and potentially introduce false positives in terms of identified residues (e.g. at position 140). Nonetheless, we identify substitutions within previously defined H9 antigenic sites at new amino acid positions and confirm them experimentally as antigenically important. This indicates that such an approach can be extended to influenza subtypes where multiple, smaller databases of antigenic data exist, such as H5 and H7 viruses. While this study used only post-infection chicken antisera for validation of antigenic residues, it is possible that antibody response induced by vaccination with inactivated, adjuvanted vaccine may differ to some degree compared with those present post-infection.

In conclusion, we show that while mAb escape mutant studies are undoubtedly helpful for understanding virus antigenicity, escape mutations often have little or no effect on inhibition by polyclonal antiserum drawn from chickens, the target species of vaccination. Additionally, many escape mutants have not been found to emerge in nature, perhaps due to negative fitness costs. Conversely, integrated modelling of sequence and HI data directly provides information on the molecular basis of antigenic variation specifically for circulating viruses, and its conclusions can be easily validated by reverse genetics. We anticipate that these results will contribute to future vaccine development and seed strain assessment by providing quantitative molecular markers that can help explain vaccine breakdown in the field and predict levels of antigenic drift during virus surveillance.

## Materials and methods

### Ethics statement

All described animal studies and procedures were carried out in strict accordance with European and United Kingdom Home Office regulations and the Animals (Scientific Procedures) Act 1986 Amendment Regulations, 2012. These studies were carried out under the United Kingdom Home Office approved project license numbers 30/2683 and 30/2952. Additionally, the work had undergone ethical scrutiny before approval by The Pirbright Institute’s Animal Welfare and Ethical Review Board (AWERB).

### Cells and eggs

MDCKs and HEK 293Ts (obtained from the European Collection of Authenticated Cell Cultures) were grown in Dulbecco modified Eagle media (DMEM) supplemented with 10% foetal bovine serum (FBS) at 37°C, 5% CO_2_. Virus was propagated in 10-day-old embryonated chicken eggs (ValoBioMedia GmbH), allantoic and amniotic fluid was harvested and pooled at 48 hours post-inoculation.

### Cloning and rescue of recombinant viruses

All viruses used in this study were generated through reverse genetics as previously described ^25,37^. Recombinant, reassortant viruses were used for generation of chicken antisera, whereby the HA of the named virus was rescued with the remaining segments from A/chicken/Pakistan/UDL-01/2008 (UDL1/08). For all other assays, high growth reassortant 1:1:6 viruses were used where the HA was taken from the named virus, the NA from UDL1/08, and the remaining 6 internal genes from A/Puerto Rico/8/1934 (PR8). Viruses with amino acid substitutions were generated through site directed mutagenesis. After propagation, all viruses were Sanger sequenced in the HA1 region to determine if any reversions/compensatory mutations had arisen, as previously described^25^.

### Antisera production

Post-infection chicken antisera were generated as described previously^38^. Briefly, 3-week-old specific pathogen free chickens were infected intranasally with 10^6^ pfu of recombinant virus. Birds were monitored twice daily for clinical signs. At 21 days post-infection birds were euthanized and bled for antisera. All other antisera were either purchased commercially or received as gifts from collaborators.

### Haemagglutination and Haemagglutination inhibition assay

Haemagglutination and HI assays were performed as described elsewhere^39^, using a solution of 1% chicken red blood cells. All HI assays with mutant viruses were performed with a minimum of three technical repeats with six individual chicken antisera. Linear mixed-effects models were used to determine whether HI titres recorded for mutant viruses were significantly different from those recorded for the parental virus. Mutant viruses were considered in a combined model with fixed effects for each combination of parental virus and for each introduced substitution. Random effects were used to account for variation in titres attributable to differences in antiserum drawn from different chickens (biological repeats) and differences between experiments (technical repeats).

Variation in HI titre associated with introduced substitutions resulting from changes in antigenicity and avidity was estimated using a similar methodology to previously described^17^. When reciprocal mutants were tested (arbitrarily termed forward and reverse for the purpose of explanation), both mutants were tested by HI against antiserum to the parental virus lacking the forward substitution (*X*) and antiserum to the parental virus lacking the reverse substitution (*Y*). The impact of the introduced substitution on antigenicity (*d*) and avidity (*v*) was estimated using:

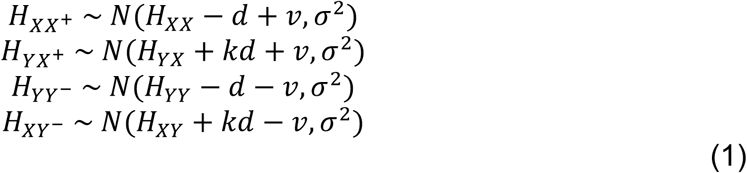

where titres for each mutant virus are assumed to be normally distributed (*N*()) with a mean depending on titres recorded for the parental virus and the antigenic and avidity effects of the introduced substitution and variance *σ^2^. H_YX_* is the log_2_ titre for the virus *X* tested with antiserum raised against the virus *Y*, and *X*^+^ and *Y*^−^ are the reciprocal mutant viruses possessing forward and reverse substitutions, respectively. Thus, *H_XY^−^_* is the log_2_ titre for the mutant virus *Y*^−^ and antiserum raised against the virus *X*. The full antigenic effect of an introduced substitution is apparent when measured using parental antiserum, but that effect may be obscured to some degree when measured using an antiserum raised against a distinct virus due to other antigenic differences. *k* is the proportion of the full antigenic impact of a substitution that is observed (*i.e*. not obscured) using antiserum raised against a distinct virus. To investigate a potential asymmetric antigenic effect, an alternative model that allowed *d* on lines one and four of Equation 1 to differ from *d* on lines two and three was used. Parameters in the above equations were co-estimated in a Bayesian model with minimally informative priors using JAGS v3.3.0 via the runjags R package v2.0.3.2 ^40,41^.

### Receptor destroying enzyme-based avidity assay

Solutions of 1% chicken red blood cells were treated with differing concentrations (8-1024 ng/ml) of a receptor destroying enzyme (RDE; neuraminidase from *Vibrio cholerae*, type II, Sigma-Aldrich cat no. N6514) for 1 hour at 37°C. Red blood cells were then washed twice and resuspended in phosphate buffered saline. 50 µL of RDE treated blood was added to 4 HAU of viruses and put on ice. Plates were read after 1 hour, the highest concentration of RDE that still allowed complete haemagglutination was recorded as the relative avidity.

### Western blotting of glycosylation mutant viruses

Allantoic fluid containing glycosylation mutants was concentrated by ultracentrifugation at 27,000 rpm for 2 hours. Concentrated viruses were then run on 7.5% SDS-PAGE gels and Western blotted. A cocktail of anti-UDL1/08 mAbs, the previously described JF7, IB3, ID2 and IG10 ^25^, were used to probe HA1, a band shift was then used to determine presence or absence of glycosylation sites.

### Genetic and antigenic data

Haemagglutination inhibition data was collated from a combination of published literature, WHO candidate vaccine virus reports, publicly available theses and previously unpublished data from our laboratory (Tables S4 and S5). All available H9N2 HA sequences were downloaded from the NCBI influenza database (https://www.ncbi.nlm.nih.gov/genomes/FLU/Database/nph-select.cgi?go=database) or the GISAID EpiFlu database (http://platform.gisaid.org). The HI dataset (see data availability statement) matched to sequence data comprised 2,547 individual titre measurements between 328 test viruses and antisera raised against 103 viruses. In total 2,131 virus-antisera combinations were represented in the dataset. The dataset included antisera drawn from various species and included both post-infection and post-vaccination (hyperimmune) antisera. 868 titres were measured using chicken hyperimmune antisera, 836 using chicken post-infection, 796 using ferret post-infection, 60 using rabbit hyperimmune, seven sheep hyperimmune, and seven goat hyperimmune.

### Phylogenetic analysis

Time-resolved phylogenetic trees were reconstructed from HA1 nucleotide sequences using a variety of models of sequence evolution implemented using BEAST v1.8.2 ^42^ and tested through comparison of Bayes factors^43^. The general time reversible model with a proportion of invariant sites and a gamma distribution describing among-site rate variation with four categories estimated from the data (GTR + I + Γ_4_) was identified as the best model of nucleotide substitution. Bayes factor analysis also determined that a codon-position model allowing rates of nucleotide substitution to vary at the third codon position, a relaxed molecular clock with branch rates drawn from a lognormal distribution^44^, and a minimally constrained Bayesian skyline demographic model should be used^45^. The maximum clade credibility tree was visualized alongside amino acid alignments using the ggtree package^46^.

### Statistical modelling and model selection

Substitutions responsible for changes in antigenic phenotype during the evolution of the virus were predicted using a modelling approach adapted from that previously described ^17^. The geometric (log_2_) HI titre was used as the response variable throughout. Mixed models constructed using the lme4 package^47^ and R v3.3.0 ^48^ including each of the following variables were assessed by likelihood ratio test: the test virus, the reference virus against which the antiserum was raised, the type of antisera used which encompasses species inoculated and method of inoculation (*i.e.* post-infection or post-vaccination), and the study from which data was collected. To prevent false support for substitutions due to repeated measurements that arise due to the evolutionary relationships between viruses, phylogenetic information was included in the model. Phylogeny branches associated with antigenicity changing events were identified as previously described^17,36,49^. Additionally, phylogenetic terms associated with changes in receptor-binding avidity and immunogenicity were identified. These branches led to clades of viruses/antisera that tended to have higher or lower titres, notwithstanding antigenic relationships. Due to the size of the search space, the optimal combination of these various binary variables reflecting phylogenetic structure was determined using random restart hill-climbing as previously described^17,50^.

To investigate the effect of amino acid substitutions at specific positions, amino acid dissimilarity between the reference virus and test virus at each variable HA1 position was tested as a predictor of reduced HI titre (p <0.05), both before and after correcting for multiple testing^51^. Positions were tested alongside phylogenetic terms, therefore substitution at an identified position must correlate with variation in titre in independent branches of the phylogeny: i.e. there are convergent, alternative, or back-substitutions associated with reduced cross-reactivity. To identify antigenically distinct amino acids at identified positions, the binary term indicating presence, or absence, of substitution was replaced with a variable with levels for every pair of amino acids observed between two viruses tested together by HI. Estimates of the antigenic impact of substitutions were made by examining associated regression coefficients for substitution terms included in combination in the absence of phylogenetic terms.

## Diversity

Diversity across different amino acid positions was calculated using Inverse Simpson’s index using all 2523 full length H9 HA sequences downloaded from the NCBI influenza database (https://www.ncbi.nlm.nih.gov/genomes/FLU/Database/nph-select.cgi?go=database) as of July 2017.

